# Genomic unveiling of the diversity in grain protein and lysine content throughout a genebank collection of winter wheat

**DOI:** 10.1101/2023.07.05.547805

**Authors:** Marcel O. Berkner, Stephan Weise, Jochen C. Reif, Albert W. Schulthess

## Abstract

Globally, wheat (*Triticum aestivum* L.) is a major source of proteins in human nutrition despite its unbalanced amino acid composition. The low lysine content in the protein fraction of wheat can lead to protein-energy-malnutrition prominently in developing countries. A promising strategy to overcome this problem is to breed varieties which combine high protein content with high lysine content. Nevertheless, this requires the incorporation of yet undefined donor genotypes into pre-breeding programs. Genebank collections are suspected to harbor the needed genetic diversity. In the 1970s, a large-scale screening of protein traits was conducted for the wheat genebank collection in Gatersleben; however, this data has been poorly mined so far. In the present study, a large historical dataset on protein content and lysine content was curated and the corresponding adjusted entry means were calculated. High-quality phenotypic data of 558 accessions was leveraged by engaging four genomic prediction approaches. Based on the predicted phenotypes of 7,651 winter wheat accessions, few of them were recommended as donor genotypes due to suitable protein characteristic. Further investigation of the passport data suggested an association of the adjusted lysine content with the elevation of the collecting site. This publicly available information can facilitate future pre-breeding activities.

**Highlight:** Historical data of lysine and protein content can be leveraged by engaging genomic prediction of an entire winter wheat genebank collection which enables to propose donor genotypes for pre-breeding.

## 1 Introduction

Worldwide, 410 Mt of consumable plant-based proteins are provided by agriculture, with soybean (*Glycine max* (L.) Merr.), maize (*Zea mays* L.), and wheat (*Triticum aestivum* L.) contributing the largest quantities (Leinonen *et al*., 2019). Unlike the first two crops, wheat is mostly used directly in human nutrition (OECD/FAO, 2021). Thus, it is not surprising that wheat provides on average 19% of the proteins consumed by humans, with some regional peaks reaching more than one third in North Africa as well as in West and Central Asia (Erenstein *et al*., 2022). Remarkable ratios were also found in some regions of South Asia: wheat consumed as flat bread accounts for about three-fifths of the daily protein consumption in Pakistani households (Hussain *et al*., 2004). Undoubtedly, the associated dominance in the diets are partially due to the prevalent cultivation in the respective regions but also wheat’s widespread availability on a global market (Shewry and Hey, 2015). Moreover, the preference for wheat can also be assigned to the specific characteristics of the protein fraction of the wheat grain which lead to the unique baking and procession quality of wheat flour (Shewry, 2009). This is one of the reasons for wheat being processed to a diversity of breads, pastries and noodles (Shewry, 2009) and as such forms a key aspect of the cuisine in many regions.

Despite the large quantity of consumed wheat protein, the nutritional quality of this protein is rather inadequate due to the unbalanced amino acid composition. In particular, shortcomings in the lysine content are the limiting factor (Leinonen *et al*., 2019) which is especially problematic since the essential amino acid lysine cannot be produced by the human organism itself and thus, must be obtained by the diet (Ufaz and Galili, 2008). On the one hand, these shortcomings can be leveled out in a diverse diet which comprises lysine-rich protein sources such as legumes, meat, fish or dairy products (Ritchie *et al*., 2018; Leinonen *et al*., 2019). On the other hand, a considerable number of people, especially in developing countries, does not have the purchasing power to diversify their diet with, for example, animal-based products (Hussain *et al*., 2004; Pellett and Ghosh, 2004; Muleya *et al*., 2022). An unbalanced wheat-rich diet may result in lysine deficiency (Meybodi *et al*., 2019). Such a deficiency is known to cause severe physical underdevelopment in children (Batool *et al*., 2015). Moreover, an inadequate supply with high quality protein can affect physiological processes, the immune system *per se* as well as the cognitive development (Batool *et al*., 2015). Impact on elderly adults is also widely reported and for this group, deficiency results in severe impairment of health including symptoms such as anemia and fatigue (Meybodi *et al*., 2019). Overall, the symptoms associated with inadequate protein supply are summarized under the name protein-energy malnutrition (Meybodi *et al*., 2019) and affect millions of people in developing countries (Batool *et al*., 2015).

Some strategies have already been proposed to increase the lysine content of staple foods. For example, artificial fortification of wheat flour with ground legumes, pseudo cereals or synthesized amino acids (Hussain *et al*., 2004) has been shown to be effective, but may have adverse effects on the processing quality or taste of end products (Meybodi *et al*., 2019). Another promising strategy could be to breed wheat varieties which combine an overall high protein content with an enrichment of lysine in the grain. In general, the potential of developing cereal crops with such characteristics has been demonstrated in maize. Naturally occurring maize mutants, such as *opaque2* and *floury2*, have been reported with a significantly elevated lysine content (Morton *et al*., 2016). In a case study, Muleya and collaborators (2022) concluded that the use of varieties with such a mutation reduces the risk of lysine deficiency by 21% for the poorest quintile of households in Malawi. To the best of our knowledge, an analog wheat variety has however not been developed so far. Lysine content has generally not been of interest in commercial wheat breeding programs and therefore, the potential of a breeding-based approach might be particularly high for such an orphan trait. Moreover, the naturally occurring lysine content of wheat grains is mainly influenced by the genotype and depends only to a small extent on environmental factors (Lawrence, 1976). Both arguments advocate for a breeding-based approach such as outlined: Firstly, the variation in lysine content of a large quantity of genotypes needs to be analyzed which is very laborious in the field and laboratory. The first step is followed by the identification of donor genotypes with a high lysine content in the protein fraction. Lastly, the favorable genetics of donor genotypes would be considered in pre-breeding activities and selectively transferred into the elite gene pool of modern breeding programs. While the latter step is mostly rather foreseeable, the first two are the bottlenecks for increasing the lysine content because they are time-consuming, demand resources and the result largely depends on the variation available for analysis.

Genebank collections for wheat are known to harbor large genetic diversity (Sansaloni *et al*., 2020; Schulthess *et al*., 2022) and phenotypic variation (Philipp *et al*., 2018; Schulthess *et al*., 2022). Thus, donor genotypes with a high content of lysine and protein could potentially be identified within these collections. Earlier attempts to screen genebank collections of wheat for both traits date back to the early 1970s. Vogel and collaborators evaluated 12,000 wheat accessions from the World Wheat Collection of the United States Department of Agriculture (USDA) (Vogel *et al*., 1976). In the same decade, both traits were measured for 9,706 *Triticum* accessions at the predecessor institution of the Federal *ex situ* Genebank of Agricultural and Horticultural Crops which is today hosted at the Leibniz Institute of Plant Genetics and Crop Plant Research in Gatersleben (IPK Genebank) (Lehmann *et al*., 1978). The aim of the aforementioned study was to screen the entire collection once and to identify accessions with a strong deviation from the population mean. The deviating accessions were re-evaluated in another year in order to account for an overestimation due to environmental effects. Until the mid-1980s, further successions were successively investigated in a structured manner (Müntz and Lehmann, 1987). Despite the sheer amount of work reflected by the work from Lehmann and collaborators (1978), this data has not been mined in depth according to today’s standards and possibilities. Since then, developments in biostatistics and genomics urge the need for a reevaluation of this historical dataset. This includes the connection of phenotypic data to genotypic data derived by next generation sequencing, which becomes more and more available for large parts of the cereal collections at the IPK Genebank (Schulthess *et al*., 2022). Combining and analyzing data will undoubtedly become more important for the work of genebank curators in the future. Since the evaluation in the 1970s, the IPK Genebank has increased in size. With more than 27 thousand genebank accessions of *Triticum* species (Oppermann, 2023), the IPK Genebank preserves nowadays the 9^th^ largest collection of plant genetic resources of wheat and its crop wild relatives (FAO, 2010). Genomic prediction could be used to characterize these new non-phenotyped parts of the collection as well as those parts without reliable phenotypic data. The power of targeted genomic prediction has recently been shown by many studies in the context of genebanks (Yu *et al*., 2016; Gonzalez *et al*., 2021; Berkner *et al*., 2022; Schulthess *et al*., 2022). Finally, informing the interested public on the newly generated information according to the FAIR (Findable, Accessible, Interoperable and Reusable) (Wilkinson *et al*., 2016) principles will further activate genebanks. This strategy could enable breeders to specifically select suitable donor genotypes and eventually, it may contribute to a future with less malnutrition in developing countries.

The main aim of this study was to activate historical records of the nutritional quality of wheat accessions stored at the IPK Genebank for their use in plant breeding and research. In more detail, we targeted (1) to curate the raw historical records for protein and lysine content which were generated between 1970 and 1986, (2) to analyze the data across years in order to generate outlier-corrected adjusted entry means for genebank accessions, (3) to apply a most suited model for genomic prediction in order to predict phenotypes for the majority of genebank accessions and (4) to suggest a set of well characterized suitable donor genotypes to breeders and the interested public.

## 2 Material and Methods

### 2.1 Curation of historical records

Historical data on protein and lysine content were compiled and curated. Some of the data originally recorded on punched tapes was unlocked; other data was recorded manually from paper files. All records were checked for accuracy and linked to the currently used accession numbers. This data originates from a large screening of the *Triticum* collection of the IPK Genebank. Between 1970 and 1986, 4,971 accessions were cultivated in 11 almost consecutive years (Fig. S1), seeds were harvested and analyzed in the laboratory for protein content and lysine content. Detailed description of the procedure has been given by Lehmann and collaborators (1978). Best linear unbiased estimates (BLUEs) for thousand grain weight (TGW) were used as published by Philipp and collaborators (2019).

### 2.2 Origin and curation of genomic data

This study relied on a genomic dataset which has been published by Schulthess and collaborators (2022). Briefly, the authors requested 7,651 accessions from the *Triticum* collection of the IPK Genebank and developed 7,745 isolate lines from them. From here onward, these isolates are referred to as accession samples. All 7,745 accession samples were genotyped by following a genotyping-by-sequencing approach. Reads were aligned to the first version of the reference genome var. Chinese Spring (IWGSC, 2018). After alignment, markers were rejected based on homozygosity of either the reference or alternative allele. In the next step, information of markers was omitted based on missing values (> 10%), a minimum homozygous allele count of < 10% and a maximum heterozygosity of > 1%. Later, imputation was done based on the dominant allele. Afterwards, further filtering based on a minor allele frequency of 1% led to a final matrix with 17,118 markers which was used for downstream analysis.

### 2.3 Outlier correction and analysis of phenotypic data

The raw data for protein and lysine content was trimmed to ensure that the data could be analyzed. Per trait, accessions were excluded from further analysis if they were represented by a single datapoint. Phenotypic values of 561 accessions remained after this trimming. Furthermore, all records of a year were omitted if no overlap with records of other years could be found. Outlier correction and calculation of BLUEs was done as described by Philipp and collaborators (2018). Briefly, the following linear mixed model was fitted to the data:

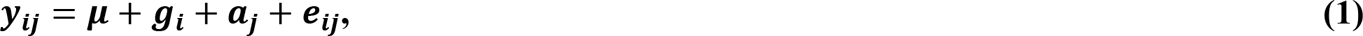

where ***y_ij_*** is the protein content (or lysine content) measured on seeds of the accession ***i*** which were harvested in the year ***j***. Accordingly, ***μ*** is the general fixed population average effect, while ***g***_***i***_ and ***a***_***j***_ represent the effects of the genotype and the year, respectively. The term ***e***_***ij***_refers to the error of the model of which the variance is modelled as specific for each year. For the identification of outliers and the estimation of BLUEs, the term ***g***_***i***_ was modelled as fixed while ***a***_***j***_was modelled as random. In contrast, both terms were considered as random for the calculation of heritability. Outliers were identified based on standardized residuals and with a correction for multiple testing (Nobre and Singer, 2011; Holm, 1979) as implemented by Philipp and collaborators (2018).

Heritabilities (***h***^**2**^) of both traits were estimated as described by Philipp and collaborators (2018),

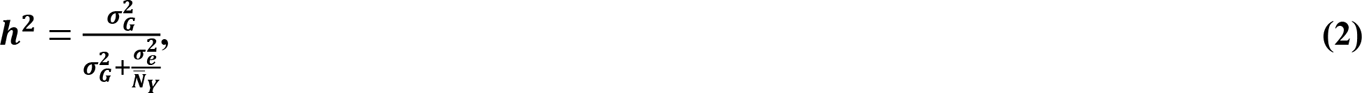

where 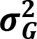 and 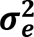 refer to the genetic variance and error variance, respectively. 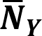 is the average number of years in which an accession was tested. In addition, above explained variances components were used to compute plot-based heritabilities as:

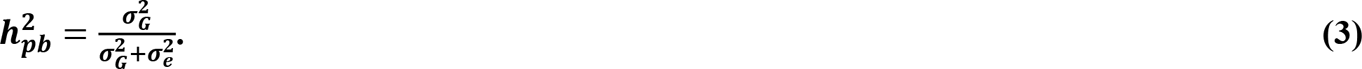

Lysine content was adjusted for protein content and TGW, because lysin content was strongly correlated with both other traits. The adjustment approach was a derivative of the approach applied by Vogel and collaborators (1975). Briefly, a multiple linear regression model was fitted on the BLUEs of lysine content in dependence on protein content and TGW. Afterwards, lysine content was adjusted genotype-wise based on the partial regression coefficients and the mean-centered protein content and TGW values as follows:

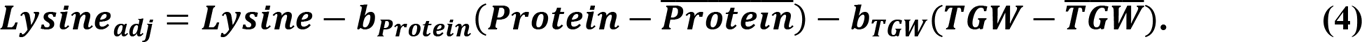

### 2.4 Analysis of population structure

The relatedness of the 7,745 genotyped accession samples was studied based on a principal coordinate (PCo) analysis (Gower, 1966). For this, pair-wise Rogers’ distances (Rogers, 1972) were calculated between genomic profiles of all accession samples and compiled into a distance matrix; the complexity of the distance matrix was reduced by deriving PCos (Gower, 1966). First and second PCos, which retain the highest amount of variation, were plotted against each other to graphically portray possible patterns resulting from population structure.

### 2.5 Genomic prediction models and their evaluation

In the present study, four different genomic prediction models, namely G-BLUP, EG-BLUP, Bayes A, and Bayesian Lasso, were compared based on their performance. The G-BLUP model (VanRaden, 2008) predicts phenotypic values based on additive genetic effects. These effects are explained by the relationship among the genotypes. The prediction model for ***n*** genotypes has the following matrix notation (Henderson, 1985):

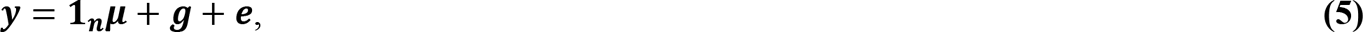

where the phenotypic values (BLUEs), given by the vector ***y***, are a function of the general mean (***μ***) and the ***n***-dimensional vectors ***g*** and ***e***, which account for the genotypic values and the model’s residuals, respectively. The ***n***-dimensional vector of ones (**1**_***n***_) assigns ***μ*** to each element of ***y***. The vectors ***g*** and ***e*** follow multivariate normal distributions 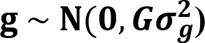 and 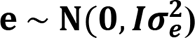 which depend on the genomic-estimated additive relationship matrix ***G*** and the genetic variance 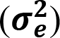 or ***I*** and the residual variance 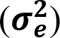, respectively. The ***n*** × ***n*** matrix ***G*** was calculated based on the first method described in VanRaden (2008) while ***I*** is an n-dimensional identity matrix.

EG-BLUP accounts for additive-by-additive epistasis (Jiang and Reif, 2015) and can be seen as an extension to the G-BLUP model as follows:

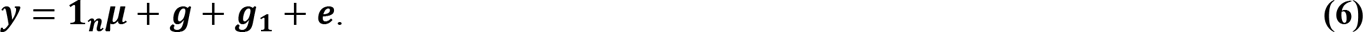

In equation **(6)**, the terms ***y***, **1**_***n***_, ***μ***, ***g***, and ***e*** are as defined in equation **(5)**. The n-dimensional vector ***g***_**1**_ accounts for the additive-by-additive effect among genotypes. This effect follows a multivariate normal distribution 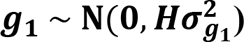 where ***H*** = ***G***#***G***, with # being the Hadamard product operator (Jiang and Reif, 2015).

BayesA (Meuwissen *et al*., 2001) was used as the base Bayesian model for genomic prediction. In this prediction approach, priors of the regression parameters are assumed to follow a scaled-t distribution. Genomic prediction with the Bayesian Lasso was applied according to Park and Casella (2008). In this approach, the regression parameters have a double-exponential prior.

The four genomic prediction models were compared based on their ability to accurately predict phenotypes. In this comparison, the unit of quality was the prediction ability, which was defined as the correlation between the BLUEs and the predicted phenotypes. The comparison was established by means of ten-fold cross-validation. All accession samples with known BLUEs were assigned to one of ten equally sized groups. Nine of these groups were incorporated in the prediction model as training set in order to predict the phenotypes in the remaining group, known as test set. The prediction was repeated in such a manner that each group has once been the test set and nine times part of the training set. Thereafter, predicted phenotypes of all test sets were combined and the Pearson correlation coefficient with the respective BLUEs was calculated. This whole process was independently repeated 100 times. For an unbiased comparison, all models were tested based on the same training and test set. The best performing prediction model was used to prediction phenotypes of all accession samples with genomic data. In the latter case, all accession samples having available phenotypic data were used as training set

All computational calculations, analysis as well as the creation of figures was implemented in the R environment (R v. 4.0.2). Solving the linear mixed model for data curation, BLUEs computation and the estimation of variance components from phenotypic data were done by engaging ASReml-R 4 v. 4.1.0.110 (Butler *et al*., 2018). Genomic prediction models were implemented with the R package BGLR v. 1.0.8 (Pérez and De Los Campos, 2014).

## 3 Results

### 3.1 Curated data with high quality

The curated phenotypic records resulted in a comprehensive dataset for protein content and lysine content which comprise 11 years of experimental trials. In total, the resulting raw dataset included 5,952 records for protein content and 5,940 records for lysine content from a total of 4,971 accessions. Across years, the raw data did not only display differences in the traits’ distributions; but moreover, the number of recorded data points differed strongly with a clear dominance for the year 1970 in which 3,442 records were taken per trait (Fig. S1). In contrast, only six measurements were reported for 1983; these were excluded due to the absent overlap with any other year. Despite the large amount of data, the dataset was rather incomplete with an unbalanced structure with most accessions tested only in one year (Table S1): Of all 4,971 accessions, 4,410 accessions were grown and characterized once without any replication. These records were excluded from further analysis to ensure that reliable BLUEs can be obtained for these accessions. After this step, remaining accessions were evaluated in up to seven years, with an average number of 2.41 and 2.39 for protein and lysine content, respectively.

The quality of the data can be reviewed based on heritability for the two traits (Table 1). For protein content, the heritability before outlier correction reached 0.77 and could only be slightly improved due to the correction. The quality of the data for lysine content improved by 23.14% due to the removal of 17 outlier data points, leading to an increase in heritability from 0.47 to 0.58. The estimated plot-based heritabilities behaved accordingly, as also evidenced by the negligible amount of rejected data points.

**Table 1.** Description of the dataset for protein content and lysine content before and after outlier correction. Depicted are the average number of datapoints per accession (***μ_N_***), the genetic variance 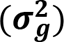, the error variance 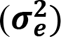 as well as the heritability (***h***^**2**^) and the plot-based heritability 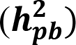. Accessions with a single datapoint per trait were disregarded here.

| Trait | Outlier correction | $\mu_N$ | $\sigma_g^2$ | $\sigma_e^2$ | $h^2$ | $h_{pb}^2$ |
| --- | --- | --- | --- | --- | --- | --- |
| Protein content | before | 2.410 | 3.894 | 2.808 | 0.77 | 0.58 |
|  | after | 2.407 | 4.017 | 2.679 | 0.78 | 0.60 |
| Lysine content | before | 2.394 | 0.087 | 0.235 | 0.47 | 0.27 |
|  | after | 2.385 | 0.096 | 0.168 | 0.58 | 0.36 |

### 3.2 Estimated average phenotypic performance of accessions and associations of the traits

The analysis resulted in BLUEs of 558 accessions for both traits, namely protein content and lysine content. On average, accessions had a protein content of 17.61% and a lysine content of 4.17‰. However, some accessions were found with a very positive deviation from the average (Fig. 1): 12 accessions exhibited a lysine content of more than 5.0‰. Protein content and lysine content were highly correlated (r = 0.63, p < 0.01). Out of the 558 accessions, only 319 accessions additionally had BLUEs for TGW. TGW was negatively correlated with both, protein content and lysine content (Fig. 1). With this information, the adjusted lysine content of these 319 accessions was calculated. The adjustment completely broke the correlations of lysine content with TGW and protein content but was still strongly correlated with lysine content *per se*.

**Fig. 1.**
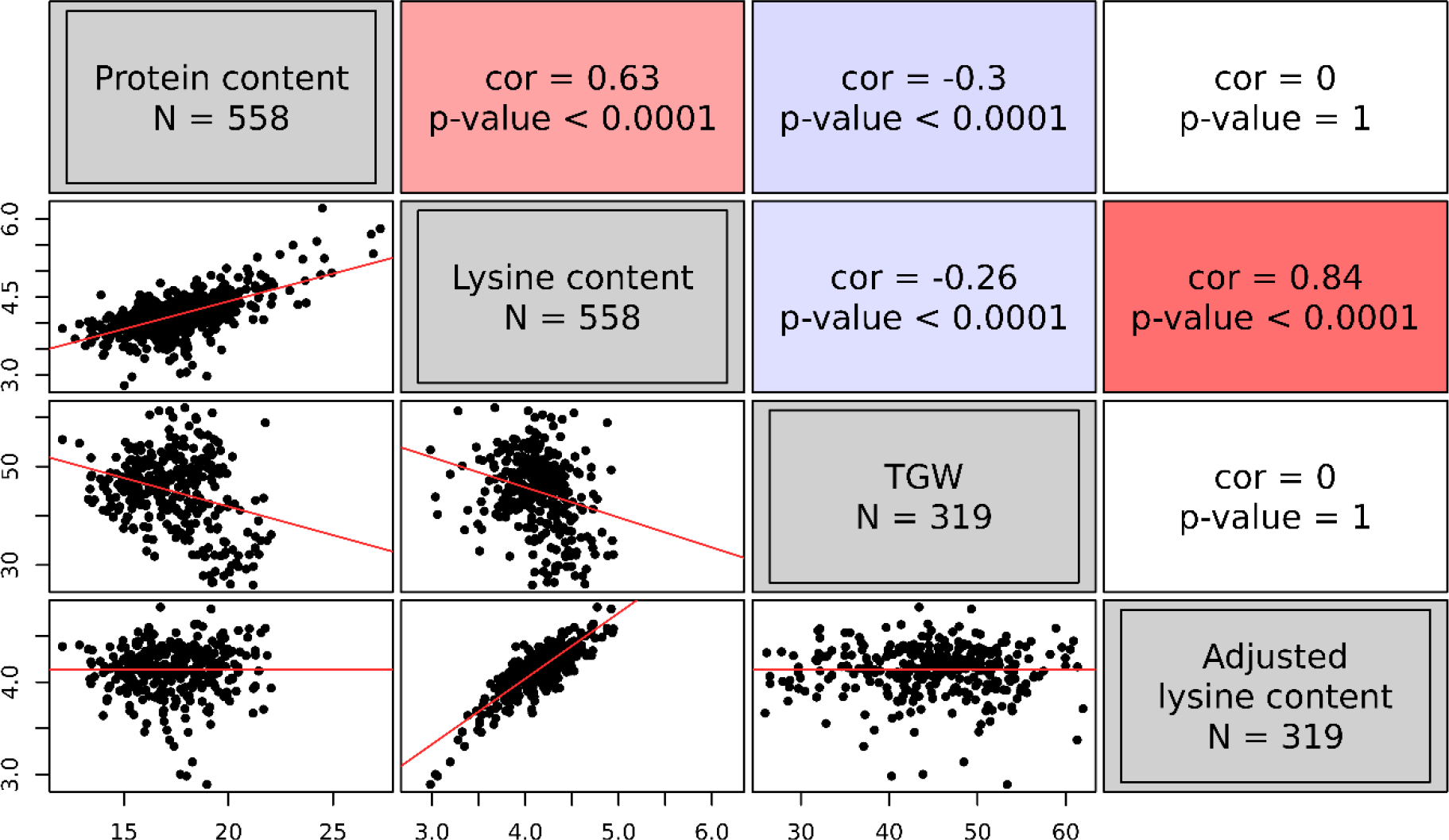
Correlations among the best linear unbiased estimates of four grain traits, namely protein content (%), lysine content (‰), thousand grain weight (TGW) (g), and adjusted lysine content (‰). The upper triangle of the correlogram depicts the Pearson correlation coefficients with the associated p-value. Positive correlations are shown in red; while, negative correlations are indicated in blue. The lower triangle of the correlogram displays the best linear unbiased estimates of the four traits plotted against each other. Regression lines are shown in red.

### 3.3 Comparison of different genomic prediction approaches

Both genotypic data and phenotypic data, which were available for 337 accession samples, were used for further genomic analysis and prediction of protein and lysine content. The number of genotyped accession samples with phenotypic information was only slightly lower for adjusted lysine content. In addition, no clear relationship pattern was observed between the distribution of the 337 accession samples along the PCos of the genomic distances and the phenotypic variation. Thus, no subpopulations were found with substantially higher or worse performing accessions compared to the population mean. All in all, the available training set with reliable phenotypes corresponds to a representative sample of the whole winter wheat collection (Fig. 2). Therefore, despite its limited size and provided high cross-validated prediction abilities, reliable predictions should be expected for both, phenotyped and non-phenotyped accessions.

**Fig. 2.**
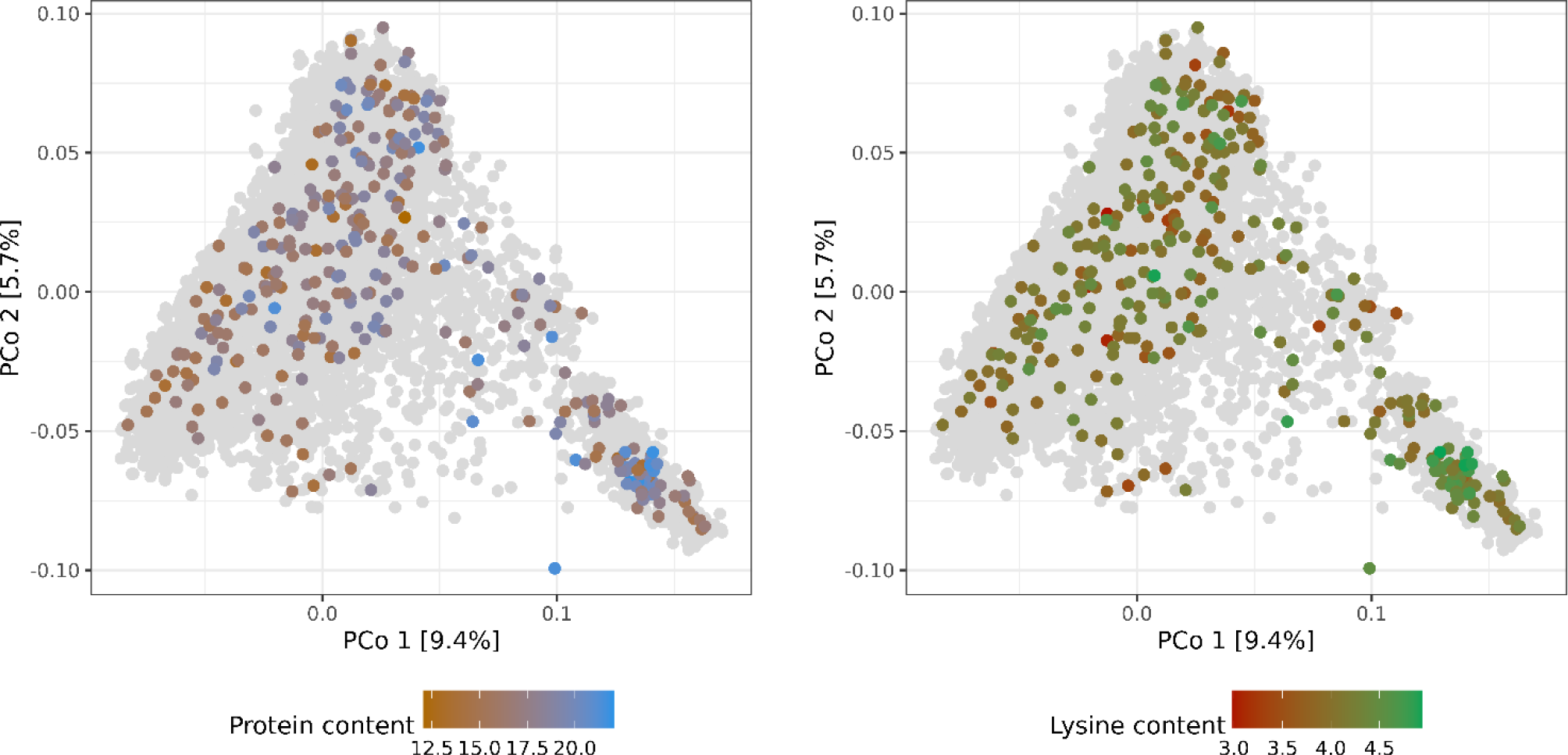
Molecular diversity of the IPK winter wheat collection covered by accession samples with best linear unbiased estimates of protein (%) and lysine (‰) content. Distributions of the 558 phenotypic values are depicted via colorcoding and shown separately per trait. Biplots are based on the first and second principal coordinates (PCo) from the Rogers’ distances between 7,745 accession samples characterized with genotyping-by-sequencing. Gray dots represent genotyped accession samples lacking phenotypic values.

Four different genome-wide prediction approaches were implemented and compared based on the correlation between BLUEs and predicted phenotypes. EG-BLUP outperformed G-BLUP and both Bayesian methods for the prediction of protein content, lysine content, and adjusted lysine content (Fig. 3). In terms of average cross-validated prediction abilities, EG-BLUP was 5.05% superior than the best of the three other alternative approaches for the prediction of adjusted lysine content. In addition, two different approaches were compared for the prediction of adjusted lysine content (Fig. S2). The separate prediction of lysine content, protein content and TGW in order to calculate the derived trait based on these predictions was marginally less accurate than using the derived trait for the genomic prediction.

**Fig. 3.**
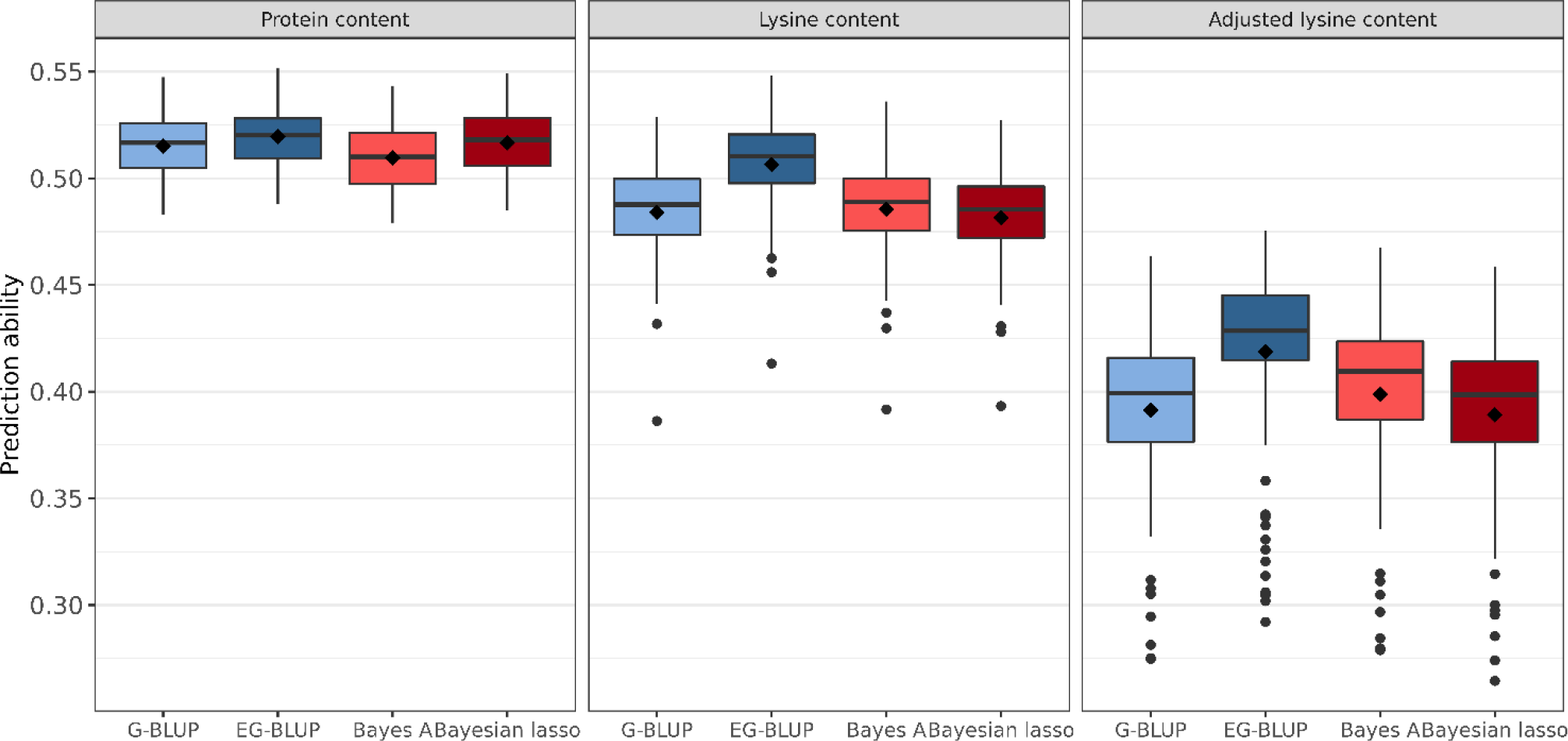
Distribution of genomic prediction abilities estimated in 100 ten-fold cross-validation runs for protein, lysine, and adjusted lysine content. Four genomic prediction models were considered: G-BLUP, EG-BLUP, Bayes A, and Bayesian lasso. Boxes enclose 50% of the central data, including median (horizontal black bold line) and mean (black diamonds), while whiskers are ±1.5× interquartile range and dots represent extreme values.

### 3.4 Predicted phenotypes of 7,745 accession samples

Protein content, lysine content, and adjusted lysine content were predicted for 7,745 accession samples by applying EG-BLUP - the most-accurate prediction model in cross-validations. For all accession samples, predicted protein content, lysine content and adjusted lysine content averaged 16.85%, 4.08‰, and 4.13‰, respectively, with associated standard deviations of 0.74%, 0.11‰, and 0.09‰, correspondingly (Fig. 4). Some accessions had outstanding values for the three traits with highest predicted values of protein content, lysine content, and adjusted lysine content amounting to 20.92%, 4.8‰, and 4.64‰, respectively. Interestingly, we found few accessions that had high values for both protein content and adjusted lysine content.

**Fig. 4.**
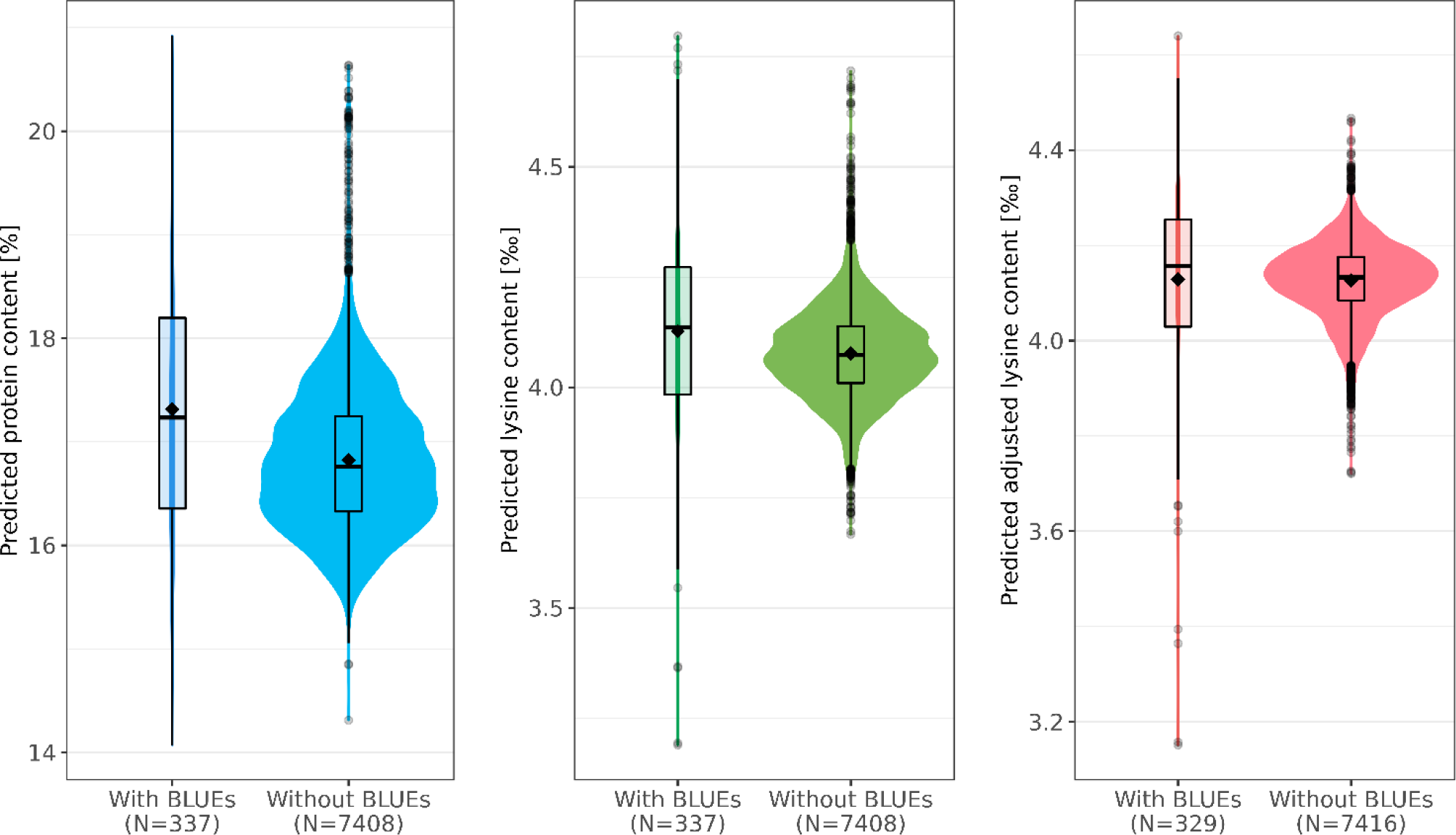
Predicted phenotypes of 7,745 wheat accession samples for protein (%), lysine(‰), and adjusted lysin (‰) content. For each trait, distributions of the prediction are shown separately for those accessions with best linear unbiased estimates (BLUEs, left) and those for which only genotypic data was present (right). Boxes enclose 50% of the central data, including median (horizontal black bold line) and mean (black diamonds), while whiskers are ±1.5× interquartile range and dots represent extreme values.

For all three traits, the size of the training set was rather small compared with the test set. For example, information of only 329 accession samples was used in order to predict the adjusted lysine content for 7,416 accession samples (Fig. 4). Interestingly, the mean and median were both lower in the test set compared with the training set. This deviation was most dominant for the predicted protein content where the means were 17.31% and 16.82% in training set and test set, respectively.

### 3.5 Definition of promising donor genotypes

The newly explored information can be used to select germplasm for pre-breeding programs with yet unexploited genetic diversity. To motivate future germplasm usage, favorable accessions were preliminary selected based on culling levels for predicted protein content and predicted adjusted lysine content in parallel. For both traits, the more stringent threshold was set to 99.9% of the normal distribution and five accessions could preliminary be selected (Fig. 5). The respective accessions had not only favorable predicted phenotypes; moreover, these accessions were also recorded with particularly high BLUEs for both traits. Thus, the prediction can be seen as a confirmation of the genetic superiority of these accessions. Additional 19 accessions were identified with a more relaxed threshold for the culling levels selection (z = 0.99) (Table S2).

**Fig. 5.**
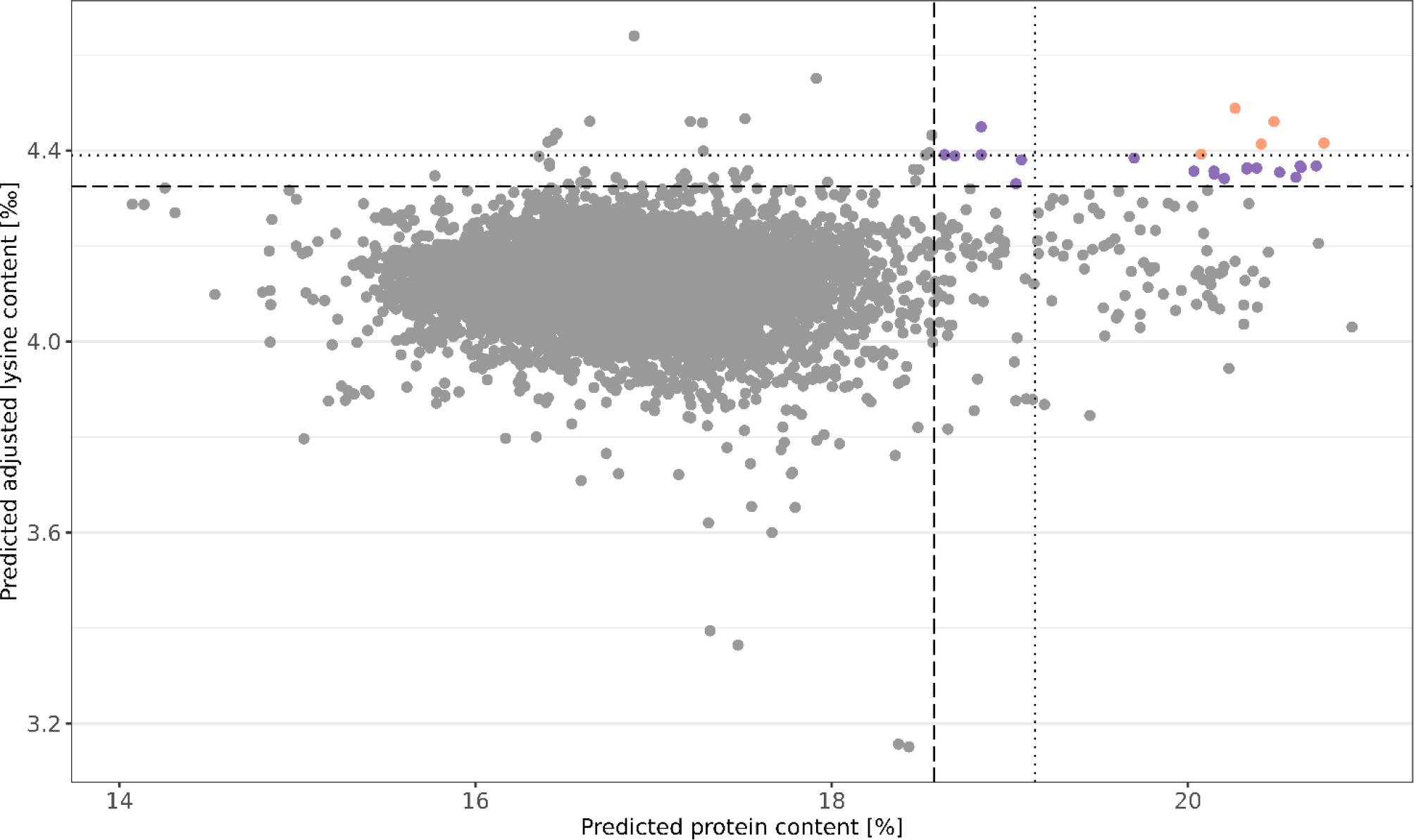
Culling levels selection based on genomic predicted protein (%) and adjusted lysin (‰) content. Shown are the predicted phenotypes of 7,745 wheat accession samples. Selection was performed with two intensities: a stringent threshold was defined by 0.999 of the normal distribution (dotted line) while a more relaxed selection threshold resulted from 0.99 of the normal distribution (dashed line). Orange and purple dots represent accessions samples which were selected based on the stringent and relaxed threshold, respectively.

### 3.6 Association of lysine with altitude

The five accessions which were preliminary selected based on the stringent culling levels selection originated from Nepal and Afghanistan (Table S2). A characteristic of these accessions is the high altitude of their collecting sites. The altitudes of the collecting sites ranged between 1,925 m and 2,975 m above sea level, as indicated by the genebank catalog (Oppermann, 2023) and the description of the collecting expedition (Witcombe, 1975). The altitude of the collecting site is only known for 927 accession samples which are 12.12% of the examined winter wheat collection. The correlation between the altitude and the predicted traits was analyzed despite the incomplete data. Predicted protein and lysine contents were positively correlated with the altitude of the collecting site; the correlation coefficient amounted to 0.23 and 0.50, respectively (p < 0.01). The predicted adjusted lysine content was also positively correlated with the altitude of the collecting site (r = 0.45, p < 0.01) (Fig8).

## 4 Discussion

The analysis of the present historical data was affected by their non-orthogonal structure. Most of the data date back to the year 1970, with the vast majority of the accessions tested without replication. Lehmann and collaborators (1978) planned to screen all accessions once with the aim of identifying accessions with a strong positive deviation from the mean of the total collection. In order to reduced cost and workload, a repetition of the analysis was conducted only for accessions with high protein content (> 18.5%) and lysin content (> 4.5‰). This strategy has two major disadvantages: First, the potential of some accessions may have been underestimated in a single testing year which would then have resulted in an erroneous rejection of this accession. Second, the total phenotypic variation cannot be estimated correctly without data from the underperforming accessions. However, precise estimates of variation are a prerequisite for optimized allocation of resources in breeding. On the one hand, this bias in the database for prediction should lower the quality of the prediction especially for the underperforming accessions (Zhao *et al*., 2012). On the other hand, this should be less relevant in our case because the selection decision in the first stage was based on data collected in one year coupled with the moderate plot-based heritabilities (Table 1). To account for the described shortcomings, we excluded accessions which were represented by only a single datapoint, as unreplicated data would provoke large uncertainty of the phenotypic data.

The exclusion of accessions with unreplicated data had strong consequences for the generated BLUEs. Trimming of the dataset reduced the number of accessions in the dataset to 11% and furthermore, resulted in higher mean values of protein content (15.92% to 17.66%) and lysine content (3.72‰ to 4.10‰). During the time of data collection, the two-step evaluation resulted in optimized selection gain and reflects the absence of computational power needed for thorough analysis. From today’s perspective, the shortcomings in the dataset highlight the need to systematically plan screenings in a way which already consider the proper statistical evaluation. Especially with limited resources, repeated phenotyping of a well-chosen subset of accessions should be favored since missing phenotypic information can be determined by genomic predictions that rely on cheap genotyping of whole genebank collections (Yu *et al*., 2016). Given the selection strategy elaborated above, it was important to investigate whether selection decisions stood in the way of a representative training population. Inspection of the PCos and distribution of phenotypic values suggests that we found this to be the case to a limited extent.

### 4.1 Strong associations between seed traits

The results showed a strong positive correlation between the BLUEs for protein and lysine content (Fig. 1). An association of these traits has already been reported based on a large screening of the USDA world wheat collection (Vogel *et al*., 1975). The authors reported an even stronger correlation of 0.804 and 0.871 for the years 1972 and 1973, respectively. Furthermore, these authors reported a slightly negative correlation of TGW with protein content (r = -0.278) and lysine content (r = -0.266), respectively, in the year 1972. Thus, these correlations are in the same order of magnitude as the correlations found in the present study. In conclusion, these correlations indicate that accessions with a higher protein content do also have a higher overall lysine content. This is hardly surprising, because lysine is part of many groups of proteins even though not in equal abundance. Furthermore, lighter grains were identified to have an overall higher protein and lysine content. Arguably, this is due to the heterogeneous distribution of both components in the wheat grain. The storage proteins in the endosperm have a significantly lower lysine content than the embryo and bran (Vogel *et al*., 1976). In line with this, the relative lysine content of wheat grains decreases during the grain filling and maturation of the seed, thus, when the endosperm increases in size (Molino *et al*., 1988). Arguably, the fraction of the endosperm on the whole grain is larger in accessions with a high TGW. This suggests that the proportion of tissues with low lysine content increases in heavier grains. On the other hand, Vogel and collaborators (1976) also found a strong correlation of 0.91 between the lysine content of the endosperm and of the whole grain and concluded that the whole grain trait values are sufficiently reliable for selection. The data set analyzed in the present study was generated based on whole grain samples (Lehmann *et al*., 1978), which thus represent a mixture of embryo, bran, and endosperm. Unfortunately, modern milling processes white flour which contains exclusively endosperm tissue (Yu and Tian, 2018). Therefore, accessions with beneficial characteristics could hypothetically rely on an elevation of the lysine content in seed tissues rarely used in human nutrition. In this regard, the distribution of amino acids should in future be further investigated especially in outperforming accessions. In the present study, the intention was to identify genotypes that have a high proportion of lysine in the protein fraction independently of the seed size; thus, the adjustment of lysine content was important to account for the described associations.

### 4.2 EG-BLUP with high potential for genomic prediction

The comparison of genomic prediction models has shown that EG-BLUP outperforms the other models in terms of prediction accuracy (Fig. 3). This is consistent with the findings of previous genomic prediction studies in wheat. EG-BLUP resulted in more accurate predictions compared with G-BLUP for the prediction of TGW, plant height, and yellow rust resistance (Berkner *et al*., 2022). The particular advantage of this model is its ability to account for additive effect but also for additive x additive epistasis (Jiang and Reif, 2015). These results highlight therefore the importance of additive x additive epistasis and are thus in line with previous finding in wheat (Jiang *et al*., 2017; Raffo *et al*., 2022). The superiority across many different traits, demonstrates the robustness of the EG-BLUP model when confronted with different genetic architectures.

According to our cross-validated comparison, genomic prediction of derived traits such as adjusted lysine content can be performed accurately (Fig. S2), but requires however some careful attention. In the present case, the implemented adjustment method relied on the availability of phenotypic values for three traits per accession. This restriction reduces the size of the training set and thus, the information which can be used for genomic prediction in a multiple-trait context (Schulthess *et al*., 2016). Moreover, the adjustment method relies on associations between lysine content and the two associated traits. These associations are, however, only valid for the examined set of genotypes and can differ between subsets of accessions such as for region-specific subpopulations. Even though the prediction of the derived trait based on the adjusted lysine content itself was most accurate (Fig. S2), it might be more appropriate to predict the basis traits separately if subsampling is planned later, if the traits are biased by subpopulations, or if the availability of data is very unbalanced across different traits.

### 4.3 Enrichment of the genebank catalog facilitates new strategies for breeding programs

The study presents three types of data, namely, curated raw data, BLUEs, and predicted phenotypes for interested stakeholders in breeding and research. Without any doubt, the estimated (BLUEs) and genomic predicted phenotypic performance can be used for targeted selection of accessions. Although all the above-mentioned data has now become publicly available, we wanted to examine specifically accessions with high protein content and adjusted lysine content. With high selection intensities (culling levels of z = 0.999), we selected five promising accessions: While one of the preliminary selected accessions came from Afghanistan, four accessions originate in the Arun valley in Nepal (Oppermann, 2023). The latter ones derived from a collecting expedition in 1971. Considering that the collecting sites of all four accessions were located in neighboring villages (Witcombe, 1975), they arguably share one common mechanism of upregulated synthesis and storage of protein and lysine. At reduced selection intensity with a culling level of z = 0.99, 19 additional accession were identified and 14 of these originate from the very same expedition to Nepal. Witcombe and Rao (1976) evaluated the accessions of that collecting journey based on 39 traits and clustered plant material based on phenotypic characteristics but ignoring the geographic proximity of the collecting sites. Interestingly, 14 accessions from the preliminary selected 18 accessions with Nepalese origin derived from the same phenotypic cluster. Witcombe and Rao (1976) identified the high altitude as one factor which leads to the common characteristics of this cluster.

### 4.4 Lysine’s suggested role in adaptation to high altitudes

Adjusted lysine content was associated with the altitude of the collecting sites of accessions. On the one hand, it could be argued that the clustering of accessions with high adjusted lysine content collected at high altitudes in Nepal was caused by a spontaneous and rare mutation unrelated to selective advantages such as adaptation to specific environmental conditions. On the other hand, high adjusted lysine content was significantly associated (r = 0.45; p < 0.01) (Fig. S3) with the altitude of the collecting site in our study for a larger sample of 927 accessions for which altitude information of the collecting site was available. This suggests that lysine content could play a role in the adaptation to high-altitude environments which share common environmental features, such as low temperatures, strong exposure to wind, drought due to lower humidity, high UV radiation, and hypoxia (Tranquillini, 1963; Jinqiu *et al*., 2021). All of these characteristics can cause abiotic stress to plants, but only the latter two are specific to high altitudes. Therefore, it could be hypothesized that high lysine content is relevant for adaptation to high ultraviolet radiation or hypoxia.

The involvement of lysine in the tolerance to various abiotic stresses, such as drought and salinity, has been summarized by Kishor and collaborators (2020). Additionally, Ding and collaborators (2016) reported a more than twofold increase in lysine content under hypoxic conditions in seedlings of rice (*Oryza sativa* L.). In contrast to this finding, the high adjusted lysine content in the present study is however not just a reaction of the wheat plant to abiotic stress. The high adjusted lysin content reflects a permanent adaptation which also results in higher lysine contents when these accessions are not facing the stresses of high altitudes. The preset study relies on field trials conducted at 110 m above sea level. Moreover, the present data reflects the lysine content of mature seeds but not of vegetative plant tissue such as seedling. For seed tissue, we can only speculate about a possible interplay of lysine with stresses such as ultra-violet radiation or hypoxia.

Abiotic stress resistance of seeds is mediated via proteins of the late embryogenesis abundant (LEA) protein families. In wheat, Liu and collaborators identified 179 genes encoding such proteins in the genome of var. Chinese Spring. These proteins cluster into eight groups with distinct characteristics. Dehydrins are one of these groups and their protective characteristics relies on the K-segment which is specifically enriched in lysine (Yang et al., 2015). In line with this, Bhattacharya and collaborators (2019) found that the amino acid composition of LEA proteins in wheat can largely rely on lysine. In the case of one analyzed protein, lysine accounted for more than one quarter of all amino acids. If abundance of such protective proteins is causal for high lysine contents remain however speculation.

The present study has not only outlined a strategy to mine historic data but also to leverage the data by genomic prediction. Moreover, this study equipped breeders and researchers with data for protein, lysine, and adjusted lysine content of in total 7,651 accession which can serve breeders to select suitable accessions for their prebreeding programs. This might build the starting point of varieties which are not just high in protein but which further have a more favorable composition of amino acids and might help to overcome protein-energy malnutrition in future.

## Abbreviations

USDA: United States Department of Agriculture
IPK: Genebank Federal ex situ Genebank of Agricultural and Horticultural Crops hosted at the Leibniz Institute of Plant Genetics and Crop Plant Research in Gatersleben
BLUEs: Best linear unbiased estimates
TGW: Thousand grain weight
LEA: Late embryogenesis abundant

## 5 Supplementary data

Table S1. Number of accessions tested in 1 to 7 years for protein content and lysine content. Table S2. Description of 24 accessions of wheat which were identified by a culling levels selection based on the predicted protein content and adjusted lysine content. Fig. S1. Distribution of protein content and lysine content separately per year of cultivation. Fig. S2. Prediction abilities for two approaches to predict the derived trait adjusted lysine content. Fig. S3. Predicted adjusted lysine content in per mille in association with the altitude of the collecting site in meter above sea level.

## 6 Tables

**Table S1.** Number of accessions tested in 1 to 7 years for protein content and lysine content.

| Trait | Number of years with records |  |  |  |  |  |  |
| --- | --- | --- | --- | --- | --- | --- | --- |
|  | 1 | 2 | 3 | 4 | 5 | 6 | 7 |
| Protein content | 4,422 | 439 | 41 | 19 | 37 | 11 | 1 |
| Lysine content | 4,423 | 439 | 43 | 21 | 36 | 8 | 1 |

**Table S2.**
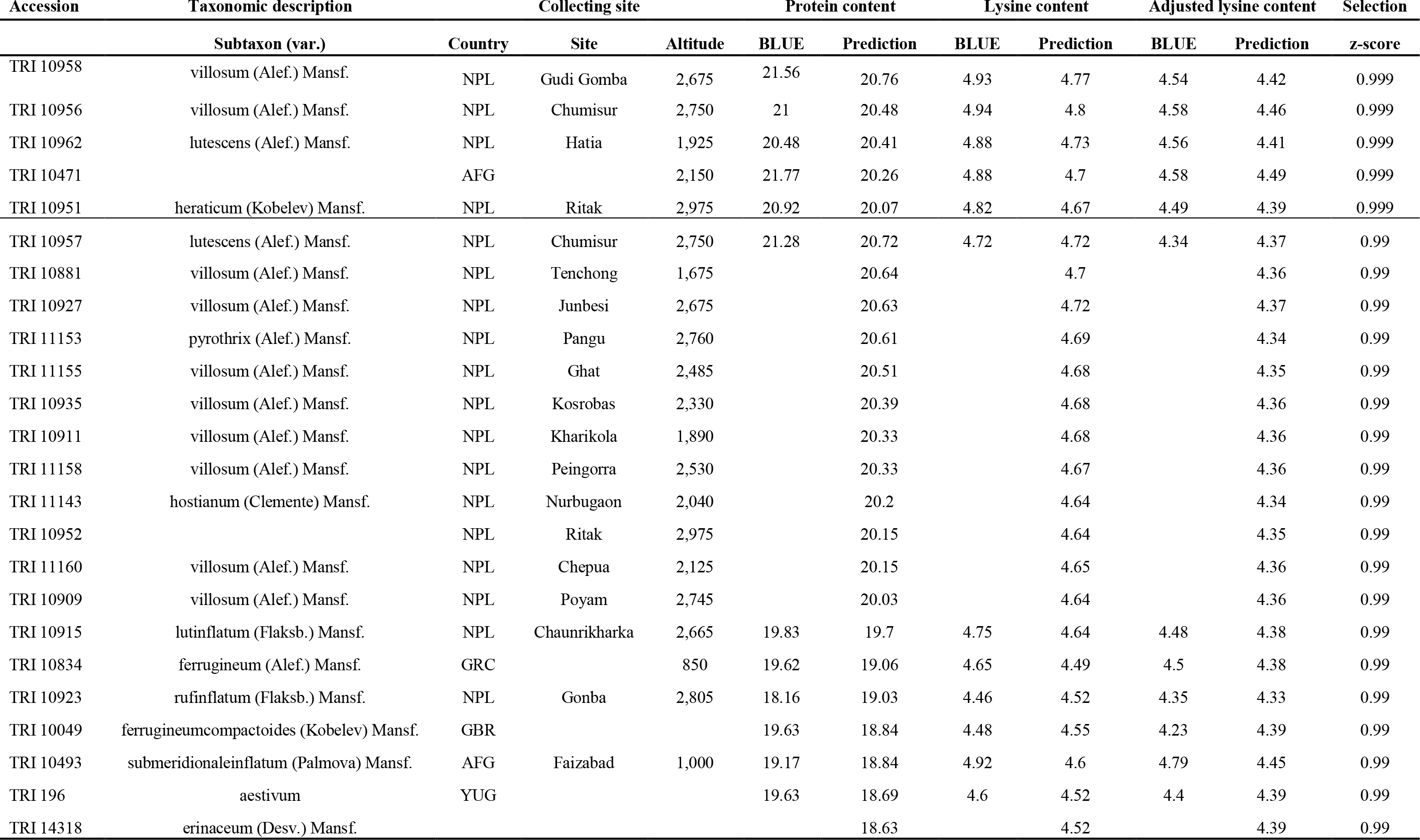
Description of 24 accessions of wheat which were identified by a culling levels selection based on the predicted protein content and adjusted lysine content. Given is the information about the taxonomic variety of *Triticum aestivum* L., the origin (Country, collecting site, altitude [m ASL]) as well as the best linear unbiased estimations (BLUEs) and predictions for protein content [%], lysine content [‰] and adjusted lysine content [‰]. Two different levels of selection intensity were appliedgiven by the z-score. Information about taxonomy and location of origin derive from Oppermann (2023) and the altitude of collecting site derives from Oppermann (2023) and Witcombe (1975).

## 7 Figures

**Fig. S1.**
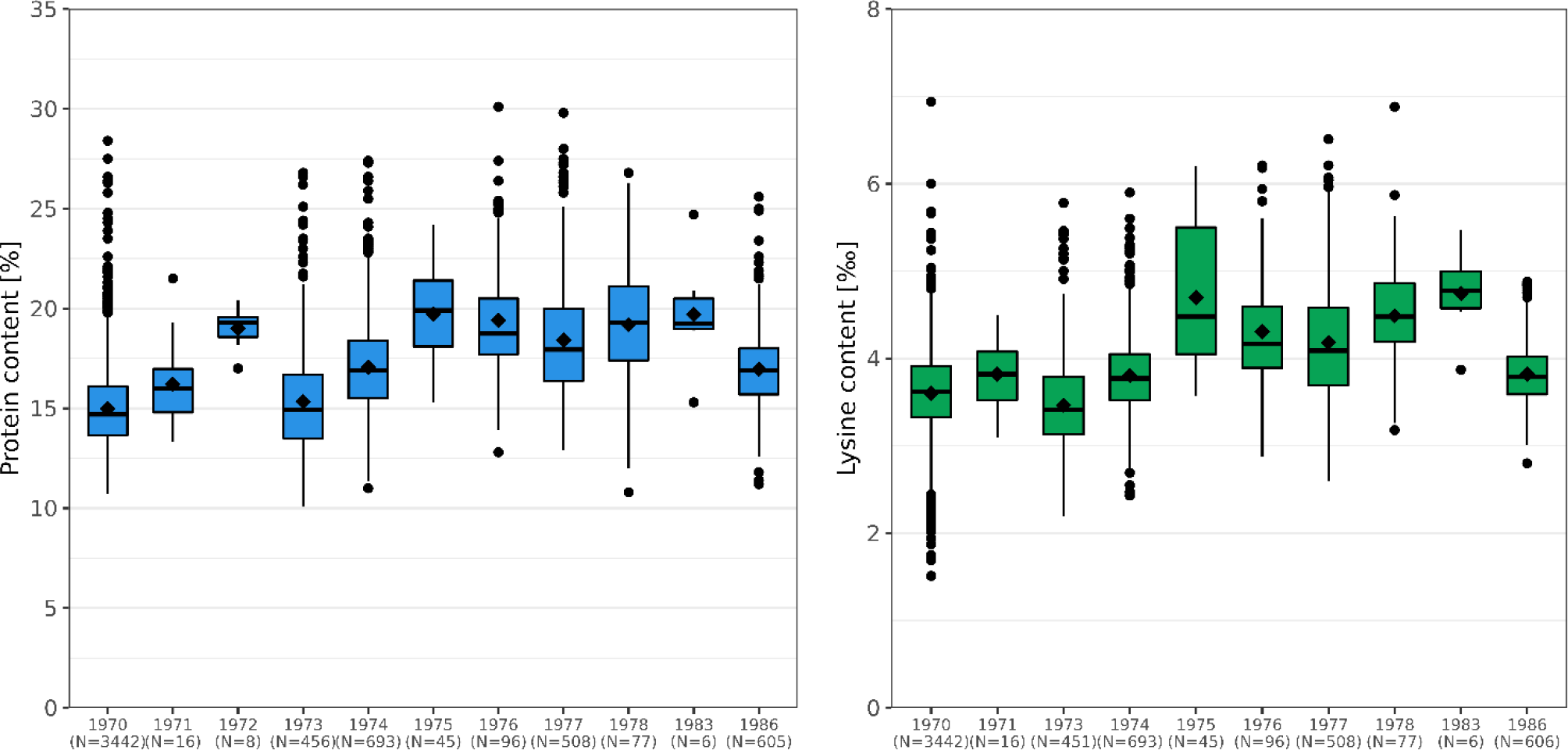
Distribution of protein content and lysine content separately per year of cultivation. Shown are the raw phenotypic values for the period 1970-1986. The number of different measurements (N) is displayed on the x-axis. Boxes enclose 50% of the central data, including median (horizontal black bold line) and mean (black diamonds), while whiskers are ±1.5× interquartile range and dots represent extreme values.

**Fig. S2.**
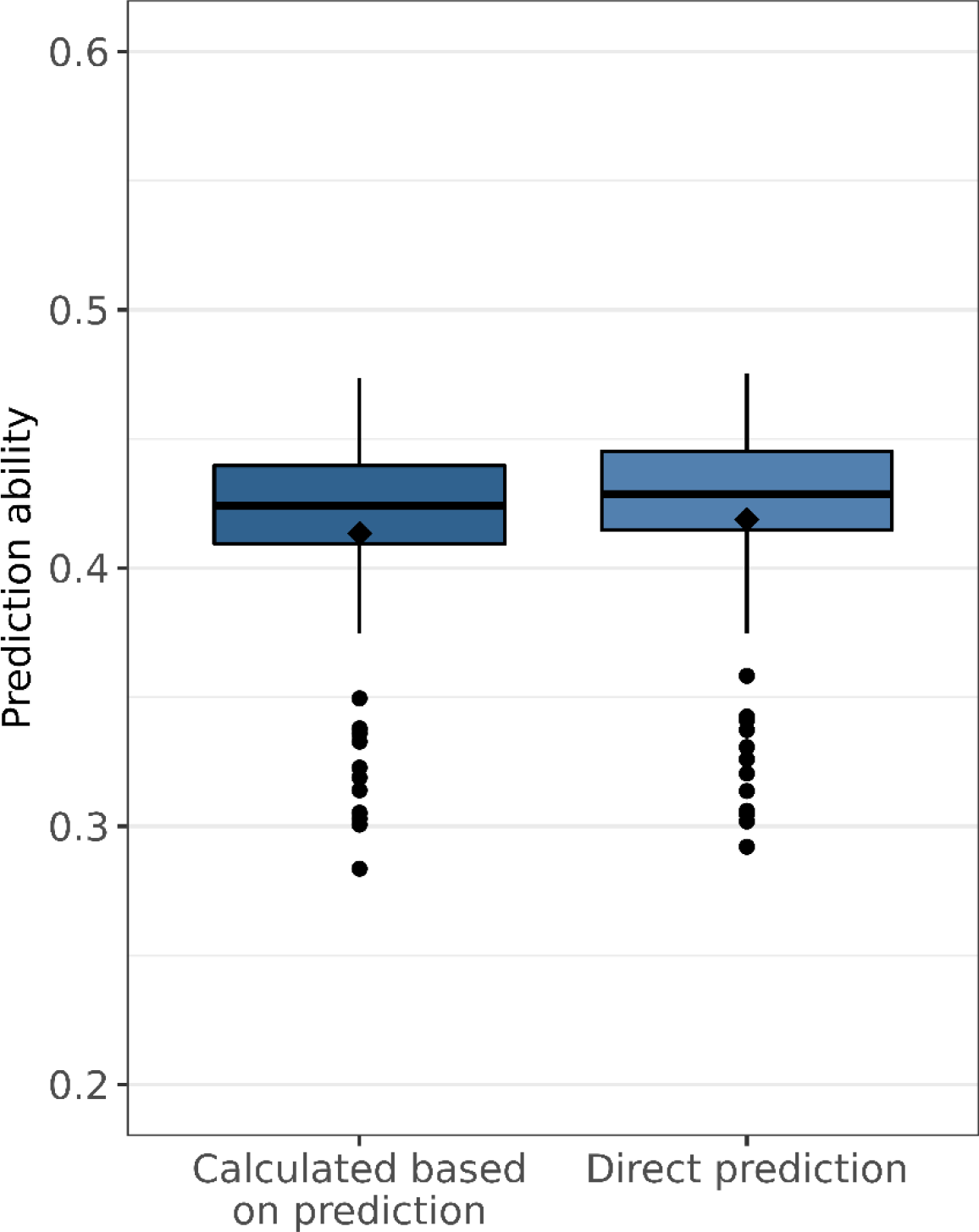
Prediction abilities for two approaches to predict the derived trait adjusted lysine content. Prediction abilities were estimated using 100 runs of ten-fold cross-validations. In the first approach, adjusted lysin content is calculated based on the predictions of lysine content, protein content, and thousand kernel weight. In the second approach, adjusted lysine content is predicted based on the adjusted lysine content calculated from the best linear unbiased estimates. Boxes enclose 50% of the central data, including median (horizontal black bold line) and mean (black diamonds), while whiskers are ±1.5× interquartile range and dots represent extreme values.

**Fig. S3.**
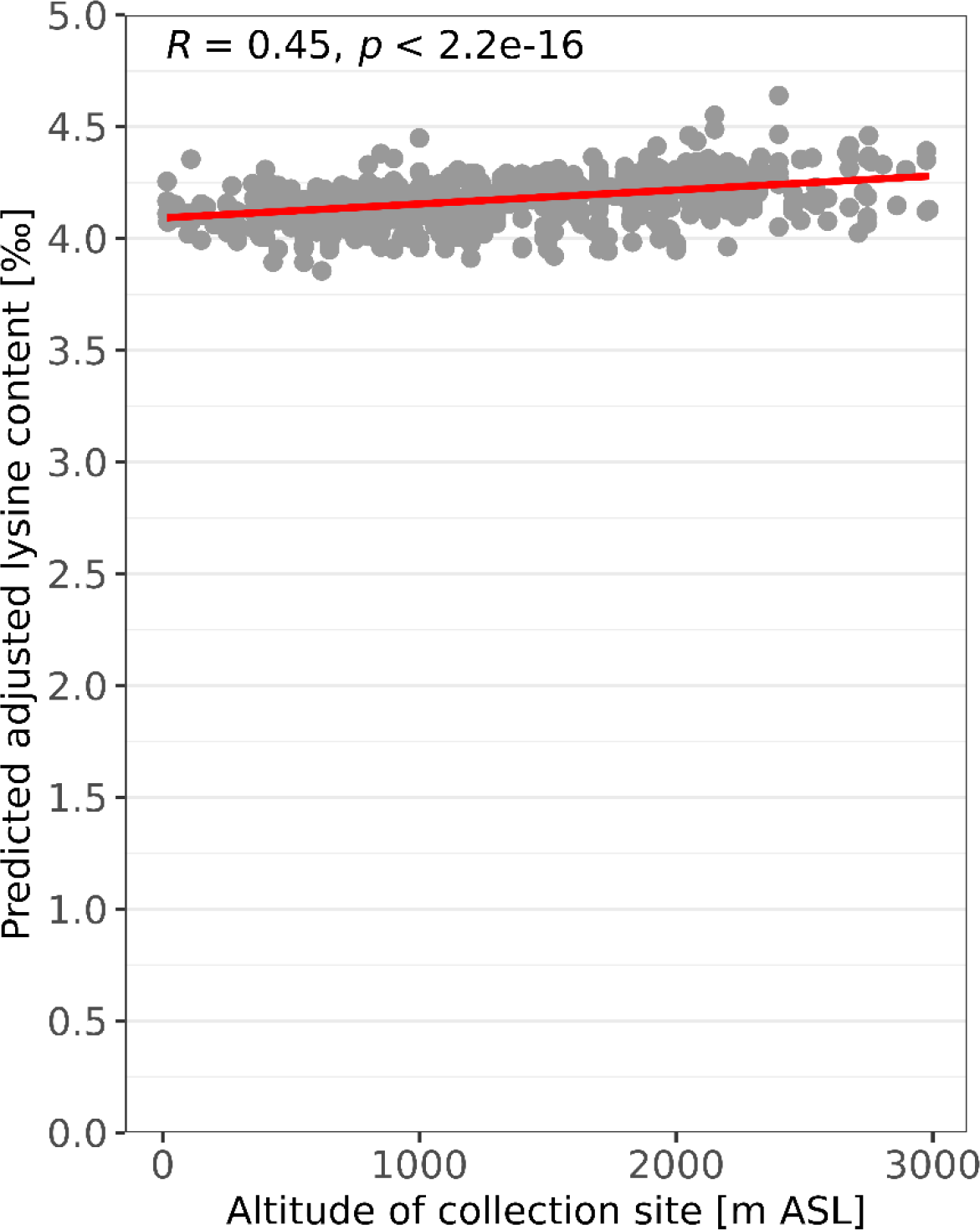
Predicted adjusted lysine content in per mille in association with the altitude of the collecting site in meter above sea level. Shown are the values of 927 accessions.

## 8 Acknowledgments

This paper is dedicated to Andreas Graner, who recognized very early the value of the historical quality data of the IPK wheat collection according to the motto “the value of a collection increases with the associated information density” and thus initiated a pillar to transform the IPK Genebank into a bio-digital resource center.

## 9 Author Contributions

SW compiled and curated the historical dataset. MOB, JCR, and AWS developed the outline of this study. MOB performed the statistical analysis with support from AWS. SW and MOB published the curated data as well as data generated in this study. MOB wrote the manuscript with support of all authors. Each author agrees that this statement reflects his contribution to this study.

## 12 Data availability

This study comprises the publication of three different type of information. These are namely the raw data in ISA-Tab format, the R code for the calculation of BLUEs and genomic prediction with all input files, and the most important output files of the analysis. The output files include BLUEs of protein and lysine content as well as the predictions of protein content, lysine content and adjusted lysine content. The aforementioned information is available via the e!DAL (Arend *et al*. 2014) online repository (DOI for e!DAL repository will be included in the galley proof of the manuscript).

